# DeepGraphMol, a multi-objective, computational strategy for generating molecules with desirable properties: a graph convolution and reinforcement learning approach

**DOI:** 10.1101/2020.05.25.114165

**Authors:** Yash Khemchandani, Stephen O’Hagan, Soumitra Samanta, Neil Swainston, Timothy J. Roberts, Danushka Bollegala, Douglas B. Kell

## Abstract

We address the problem of generating novel molecules with desired interaction properties as a multi-objective optimization problem. Interaction binding models are learned from binding data using graph convolution networks (GCNs). Since the experimentally obtained property scores are recognised as having potentially gross errors, we adopted a robust loss for the model. Combinations of these terms, including drug likeness and synthetic accessibility, are then optimized using reinforcement learning based on a graph convolution policy approach. Some of the molecules generated, while legitimate chemically, can have excellent drug-likeness scores but appear unusual. We provide an example based on the binding potency of small molecules to dopamine transporters. We extend our method successfully to use a multi-objective reward function, in this case for generating novel molecules that bind with dopamine transporters but not with those for norepinephrine. Our method should be generally applicable to the generation *in silico* of molecules with desirable properties.

## 1 Introduction

The *in silico* (and experimental) generation of molecules or materials with desirable properties is an area of immense current interest (e.g. [1-28]). However, difficulties in producing novel molecules by current generative methods arise because of the discrete nature of chemical space, as well as the large number of molecules [29]. For example, the number of drug-like molecules has been estimated to be between 10^23^ and 10^60^ [30-34]. Moreover, a slight change in molecular structure can lead to a drastic change in a molecular property such as binding potency (so-called activity cliffs [35-37]).

Earlier approaches to understanding the relationship between molecular structure and properties used methods such as random forests [38, 39], shallow neural networks [40, 41], Support Vector Machines [42], and Genetic Programming [43]. However, with the recent developments in Deep Learning [44, 45], deep neural networks have come to the fore for property prediction tasks [3, 46-48]. Notably, Coley *et al*. [49]] used Graph convolutional networks effectively as a feature encoder for input to the neural network.

In the past few years, there have been many approaches to applying Deep Learning for molecule generation. Most papers use the Simplified Molecular-Input Line-Entry System (SMILES) strings as inputs [50], and many use a Variational AutoEncoder architecture (e.g. [3, 17, 51]), with Bayesian Optimization in the latent space to generate novel molecules. However, the use of a sequence-based representational model has a specific difficulty, as any method using them has to learn the inherent rules, in this case of SMILES strings. More recent approaches, such as Grammar Variational AutoEncoders [52, 53] have been developed in attempts to overcome this problem but still the molecules generated are not always valid. Some other approaches try to use Reinforcement Learning for generating optimized molecule [54]. However, they too make use of SMILES strings which as indicated poses a significant problem. In particular, the SMILES grammar is entirely context-sensitive: the addition of an extra atom or bracket can change the structure of the encoded molecule dramatically, and not just ‘locally’ [55].

Earlier approaches have tended to choose a specific encoding for the molecules to be used as an input to the model, such as one hot encoding [56, 57], Extended Connectivity Fingerprints [58, 59] and SMILES strings directly. We note that these encodings do not necessarily capture the features that need to be obtained for prediction of a specific property (and all encodings extract quite different and orthogonal features [60]).

In contrast, the most recent state-of-the-art methods, including hypergraph grammars [61], Junction Tree Variational Auto Encoders [62] and Graph Convolutional Policy Networks [34], use a graphical representation of molecules rather than SMILES strings and have achieved 100% validity in molecular generation. Graph-based methods can be seen as a more natural representation of molecules as substructures map directly to subgraphs, but subsequences are usually meaningless. However, these have only been used to compare the models on deterministic properties such as the Quantitative Estimate of Drug-likeness (QED) [63], logP, etc. that can be calculated directly from molecular structures (e.g. Using RDKit, http://www.rdkit.org/). For many other applications, molecules having a higher score for a specific *measured* property are more useful. We here try to tackle this problem.

## 2. Methods

Our system consists of two parts: Property Prediction and Molecular Generation. For both the parts, we represent the molecules as graphs [64] since they are a more natural representation than are SMILES strings, and substructures are simply subgraphs. We train a model to predict the property scores of the molecules, specifically the binding constant of various molecules at the dopamine and norepinephrine transporters (using a dataset from BindingDB). The first part, used for (training) the property prediction part, is a Graph Convolutional Network as a feature encoder together with a Feed Forward Network. We also use an Adaptive Robust Loss Function (as suggested by [65]) since the experimental data are bound to be error prone. For the Molecular Generation task, we use the method proposed by You and colleagues [34]. In particular, we (and they) use Reinforcement Learning for this task since it allows us to incorporate both the molecular constraints and the desired properties using reward functions. This part uses graph convolution policy networks (GCPNs), a model consisting of a GCN that predicts the next action (policy) given the molecule state. It is further guided by expert pretraining and adversarial loss for generating valid molecules. Our code (https://github.com/dbkgroup/prop_gen) is essentially an integration of the property prediction code of Yang and colleagues [66, 67] (https://github.com/swansonk14/chemprop) and the reinforcement learning code provided by You and colleagues [34].

### 2.1 Molecular Property Prediction

As noted, the supervised property prediction model consists of a graph-convolution network for feature extraction followed by a fully interconnected feedforward network for property prediction.

#### 2.1.1 Feature Extraction

We represent the molecules as directed graphs, with each atom (*i*) having a feature vector *F*_*i*_ (ℝ^133^) and each bond (between atom *i* & *j*) having feature vector *F*_*ij*_ (ℝ^14^). For each incoming bond a feature vector is obtained by concatenating the feature vector of the atom to which the bond is incoming and the feature vector of the bond. Thus the input tensor is of the size *N*_bonds_ x ℝ^147^. The Graph Convolution approach allows the message (feature vector) for a bond to be passed around the entire graph using the approach described below.

The initial atom-bond feature vector that we use incorporates important molecular information that the GCN encoder can then incorporate in later layers. The initial representations for the atom and bond features are taken from https://github.com/swansonk14/chemprop and summarized in Table 1, below. Each descriptor is a one-hot vector covering the index-range represented by it (except the Atomic Mass). For Atomic Number, Degree, Formal Charge, Chiral Tag, Number of Hydrogens and Hybridization, the feature vector contains one additional dimension to allow uncommon values (values not in the specified range).

**Table 1.** Atom and bond features used in the present work

| INDICES | ATOM DESCRIPTION |
| --- | --- |
| 0 – 100 | Atomic Number (1 to 100) |
| 101-107 | Degree (1 to 5) |
| 108-113 | Formal Charge (-2 to + 2) |
| 114-118 | Chiral Tag (0 to 4) |
| 119-124 | Number of Hydrogens (0 to 4) |
| 125-130 | Hybridization (SP, SP2, SP3, SP3D, SP3D2) |
| 131 | Aromatic Atom |
| 132 | Atomic Mass * 0.01 |
| INDICES | BOND DESCRIPTION |
| 133 | Bond Present |
| 134-136 | Bond Type (Single, Double, Triple) |
| 137 | Aromatic Bond |
| 138 | Conjugated Bond |
| 139 | Bond present in Ring |
| 140-146 | Bond Stereo Code (RdKit) |

The initial atom-bond feature vector is then passed through a linear layer followed by ReLU Activation [68, 69] to get the Depth-0 message vector for each bond. For each bond, the message vectors for the neighbouring bonds are summed up (Convolution step) and passed through a linear layer followed by ReLU and a Dropout layer to get the Depth-1 message vectors. This process is continued up to a specified Depth-(N-1) message vectors. To get the Depth-N message vectors, the Depth-(N-1) vectors of all the incoming bonds for an atom are summed and then passed through a dense layer followed by ReLU and Dropout. The final graph embedding for the molecule is obtained by averaging the depth-N message vectors over all the atoms. The exact details for this model can be found in section 3.1.1.

#### 2.1.2 Regression

To perform property prediction the embedding extracted by the GCN is fed into a fully connected network. Each intermediate layer consists of a Linear Layer followed by ReLU activation and Dropout that map the hidden vector to another vector of the same size. Finally the penultimate nodes are passed through a Linear Layer to output the predicted property score. The K_i_ values present in the dataset were obtained experimentally so might contain experimental errors. If we were to train our model with a simple loss function such as root mean square (RMS) error loss, it would not be able to generalize well because of the presence of outliers in the training set. Overcoming this problem requires training the data with the help of a robust loss function that takes care of the outliers present in the training data. There are several types of robust loss functions such as Pseudo-Huber loss [70], Cauchy loss, etc., but each of them has an additional hyperparameter value (for example δ in Huber Loss) which is treated as a constant while training. This means that we have to manually tune the hyperparameter each time we train to get the optimum value which may result in extensive training time. To overcome this problem, as proposed by [65], we have used a general robust loss function that has the hyperparameters as shape parameter (α) which controls the robustness of the loss, and the scale parameter (c) which controls the size of the loss’s quadratic bowl near x=0. This loss is dubbed as a “general” loss since it takes the form of other loss functions for particular values of α. (e.g L2 loss for α=2, Charbonnier loss for α=1, Cauchy loss for α=0). The authors also propose that “by viewing the loss function as the negative log likelihood of a probability distribution, and by treating robustness of the distribution as a latent variable” we can use gradient-based methods to maximize the likelihood without manual parameter tuning. In other words, we can now train the hyperparameters α and c rather which overcomes the earlier problem of manually tuning the hyperparameters. The loss function and the corresponding probability distribution are described in Eq. 1 and Eq. 2 respectively.

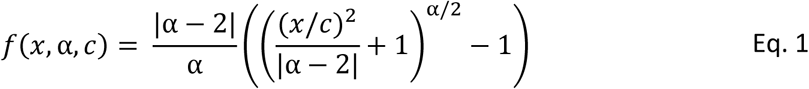

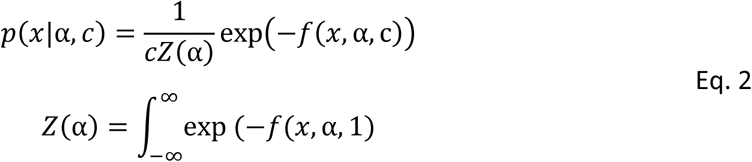

### 2.2 Reinforcement Learning for Molecular Generation

#### 2.2.1 Molecular Representation

We follow the method described by the GCPN paper [34] for the molecular generation task, with the difference being that the final property reward is the value calculated by the previously trained model for the newly generated molecules. As in the previous part, we represent the molecules as graphs, more specifically as (*A, E, F*) where *A* ∈ {0, 1}^*n*×*n*^ is the adjacency matrix, *F* ∈ ℝ^*n*×*d*^ is the node (atom) feature matrix and *E* ∈ {0, 1}^3×*n*×*n*^ is the edge-conditioned adjacency tensor (since the number of bond-types is 3, namely single, double and triple bond), with *n* being the number of atoms and *d* being the length of feature vector for each atom. More specifically, *E*_*i,j,k*_ *=* 1 if there exists a bond of type *i* between atoms *j* and *k*, and *A*_*j,k*_ *=* 1 if there exists any bond between atoms *j* and *k*.

#### 2.2.2 Reinforcement Learning setup

Our model environment builds a molecule step by step with the addition of a new bond in each step. We treat graph generation as a Markov Decision Process such that the next action is predicted based only on the current state of the molecule, not on the path that the generative process has taken. This reduces the need for sequential models such as RNNs and the disadvantages of vanishing gradients associated with them, as well as reducing the memory load on the model. More specifically, the decision process follows the equation: *p (s*_*t*+1_ |*s*_*t*_, … *s*_0_) *= p(s*_*t*+1_|*s*_*t*_), where p is the probability of the next state (*s*_*t*+1_) given the previous state (*s*_*t*_).

We can initialize the generative process with either a single C atom (as in Experiments 1 and 2) or with another molecule (as in Experiments 3, 4 and 5). At any point in the generation process, the state of the environment is the graph of the current molecule that has been built up so far. The action space is a vector of length 4 which contains the information – First Atom, Second Atom, Bond type and Stop. The stop signal is either 0 or 1 indicating whether the generation is complete, based on valence rules. If the action defies the rules of chemistry in the resultant molecule, the action is not considered and the state remains as it is.

We make use of both intermediate and final rewards to guide the decision-making process. The intermediate rewards include stepwise validity checks such that a small constant value is added to the reward if the molecule passes the valency checks. The final reward includes the pK_i_ value of the final molecule as predicted by the trained model and the validity rewards (+1 for not having any steric strain and +1 for absence of functional groups that violate ZINC functional group filters). Two other metrics are the quantitative estimation of drug-likeness (QED) [63] and the synthetic accessibility (SA) [71] score. Since our final goal is to generate drug-like molecules that can be synthetically generated, we also add the QED and 2*SA score of the final molecule to the reward.

Apart from this, we also use adversarial rewards so that the generated molecules resemble (prediction) the given set of molecules (real). We define the adversarial rewards *V*(*π*_*θ*_, *D*_Φ_) in Eq 3.

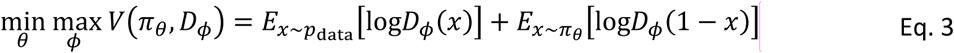

where *π*_θ_ is the policy network, *D*_*φ*_ is the discriminator network, *x* represents the input graph and *p*_data_ is the underlying data distribution which is defined either over final graphs (for final rewards) or intermediate graphs (for intermediate rewards) (just as proposed by You and colleagues [34]). Alternate training of generator (policy network) and discriminator by gradient descent methods will not work in our case since *x* is a non-differentiable graph object. Therefore we add – *V*(*π*_*θ*_, *D*_Φ_) to our rewards and use policy gradient methods [72] to optimize the total rewards. The discriminator network comprises a Graph Convolutional Network for generating the node embedding and a Feed Forward Network to output whether the molecule is real or fake. The GCN mechanism is same as that of the policy network which is described in the next section.

#### 2.2.3 Graph Convolutional Policy Network

We use Graph Convolutional Networks (GCNs) as the policy function for the bond prediction task. This variant of graph convolution performs message passing over each edge type for a fixed depth L. “The node embedding for the next depth (*l* + 1) is calculated as described in Eq. 4

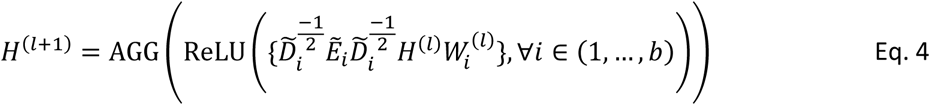

where E_i_ is the *i*^*th*^ slice of the tensor *E*, 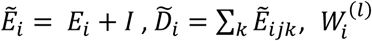 is a trainable weight matrix for the *i*^*th*^ edge type, and *H*^*(l*)^ is the node embedding learned in the *l*^*th*^ layer with *H*^*(l*)^ ∈ ℝ^*(n*+*c*)×*d*^ [34]. n is the number of atoms in the current molecule and c is the number of possible atom types (C,N,O etc.) that can be added to the molecule (one atom is added in each step) with *d* representing the dimension of the embedding. We use mean over the edge features as the Aggregate (AGG) function to obtain the node embedding for a layer. This process is repeated *L* times until we get the final node embedding.

This node embedding X is then used as the input to four Multilayer Perceptrons (MLP, denoted by m), that map a matrix *Z* ∈ ℝ^*p*×*d*^ to ℝ^*p*^ representing the probability of selecting a particular entity from the given p entities. The specific entity is then sampled from the probability distribution thus obtained. Note that since the action space is a vector of length 4, we use 4 perceptrons to sample each component of the vector. The first atom has to be from the current molecule state while the second atom can be from the current molecule (forming a cycle) or a new atom outside the molecule (adding a new atom). For selecting the first atom, the original embedding X is passed to the MLP *m*_*f*_ and outputs a vector of length equal to n. While selecting the second atom, the embedding of the first atom 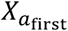 is concatenated to the original embedding X and passed to the MLP *m*_*s*_ giving a vector of length equal to *n* + *c*. While selecting the edge type, the concatenated embedding of the first 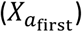 and second 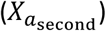 atom is used as an input to MLP m_e_ and outputs a vector of length equal to 3 (number of bond types). Finally, the mean embedding of the atoms is passed to MLP m_t_ to output a vector of length 2 indicating whether to stop the generation. This process is described in equations 5-9.

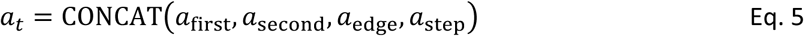

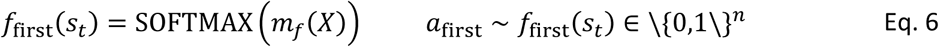

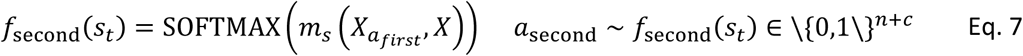

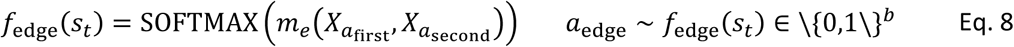

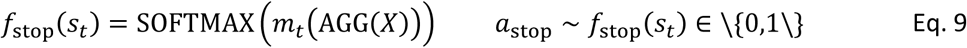

#### 2.2.4 Policy Gradient Training

For our experiments, we use Proximal Policy Optimization (PPO) [72], the state-of-the-art policy gradient method, for optimizing the total reward. The objective function for PPO is described in Eq 10.

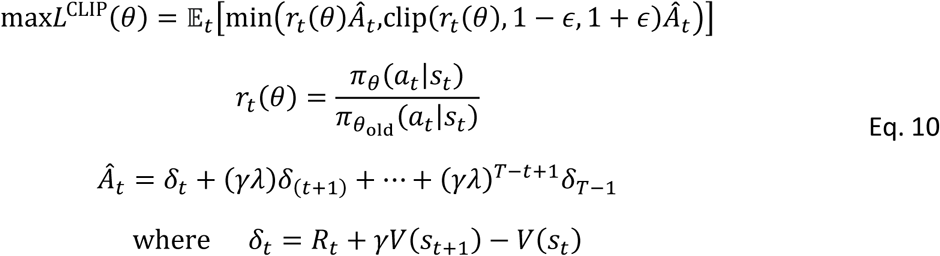

Here *s*_*t*_, *a*_*t*_, *R*_*t*_ are the state, action and reward respectively at timestep *t, V(s*_*t*_) is the value associated with state *s*_*t*_, *π*_θ_ is the policy function and *γ* is the discount factor. Also note that *Â*_*t*_, which is an estimator of the advantage function at timestep *t*, has been estimated using Generalized Advantage Estimation [73] with the GAE parameter *λ*, since it reduces the variance of the estimate.

For estimating the value of V we use an MLP with the embedding X as the input. Apart from this, we also use expert pretraining [74] which has shown to stabilise the training process. For our experiment, any ground truth molecule can be used as an expert for imitation. We randomly select a subgraph *Ĝ* from the ground truth molecule *Ĝ* as the state. *ŝ*_*t*_ The action *â*_*t*_ is also chosen randomly such that it adds an atom or bond in the graph *Ĝ\ Ĝ*. This pair *(ŝ*_*t*,_ *â*_*t*_) is used for calculating the expert loss.

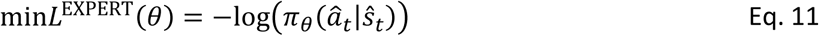

Note that we use the same dataset of ground truth molecules for calculating the expert loss and the adversarial rewards. For the rest of the paper, we will call this dataset the “expert dataset” and the random molecule selected from the dataset the “expert molecule”.

## 3. System evaluation

In this section we evaluate the system described above on the task of generating small molecules that interact with the dopamine transporter but not (so far as possible) with the norepinephrine transporter.

### 3.1 Property Prediction

In this section we evaluate the performance of the supervised property prediction component. Dopamine Transporter binding data was obtained from www.bindingdb.org (https://bit.ly/2YACT5u). The training data consist of some molecules which are labelled with their K_i_ values and some which are labelled with IC_50_ values. For this paper, we have used IC_50_ values and K_i_ values interchangeably in order to increase the size of the training dataset. Molecules having large K_i_ values in the dataset were not labelled accurately (with labels such as ∼1000) but the use of a robust loss function allowed us to incorporate these values directly. As stated above we use log transformed values (pKi). (We also attempted to learn the K_i_ values of the molecules, but the distribution was found to be heteroscedastic; hence we focus on predicting the pK_i_ values.) Data are shown in Fig 4A for the dopamine transporter and 4B for the norepinephrine transporter pKi values.

**Figure 1.**
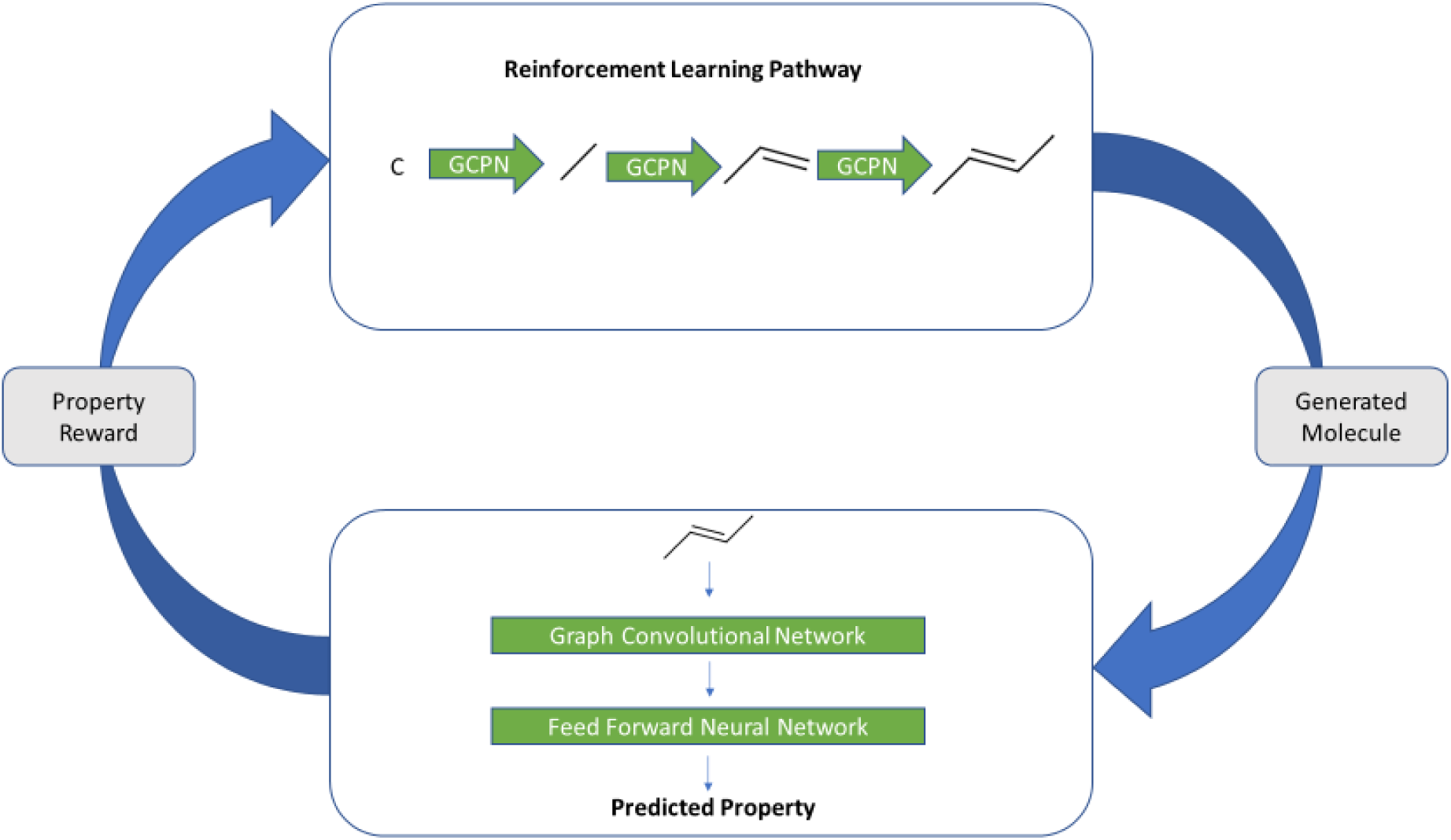
Block diagram of our basic system. A molecule is generated by the Reinforcement Learning (RL) pathway using a Graph Convolutional Policy Networks. This molecule is then used as an input for the property prediction module which outputs the property score as predicted by the module. This score is then used as the reward feedback for the RL pathway and the cycle restarts

**Figure 2.**
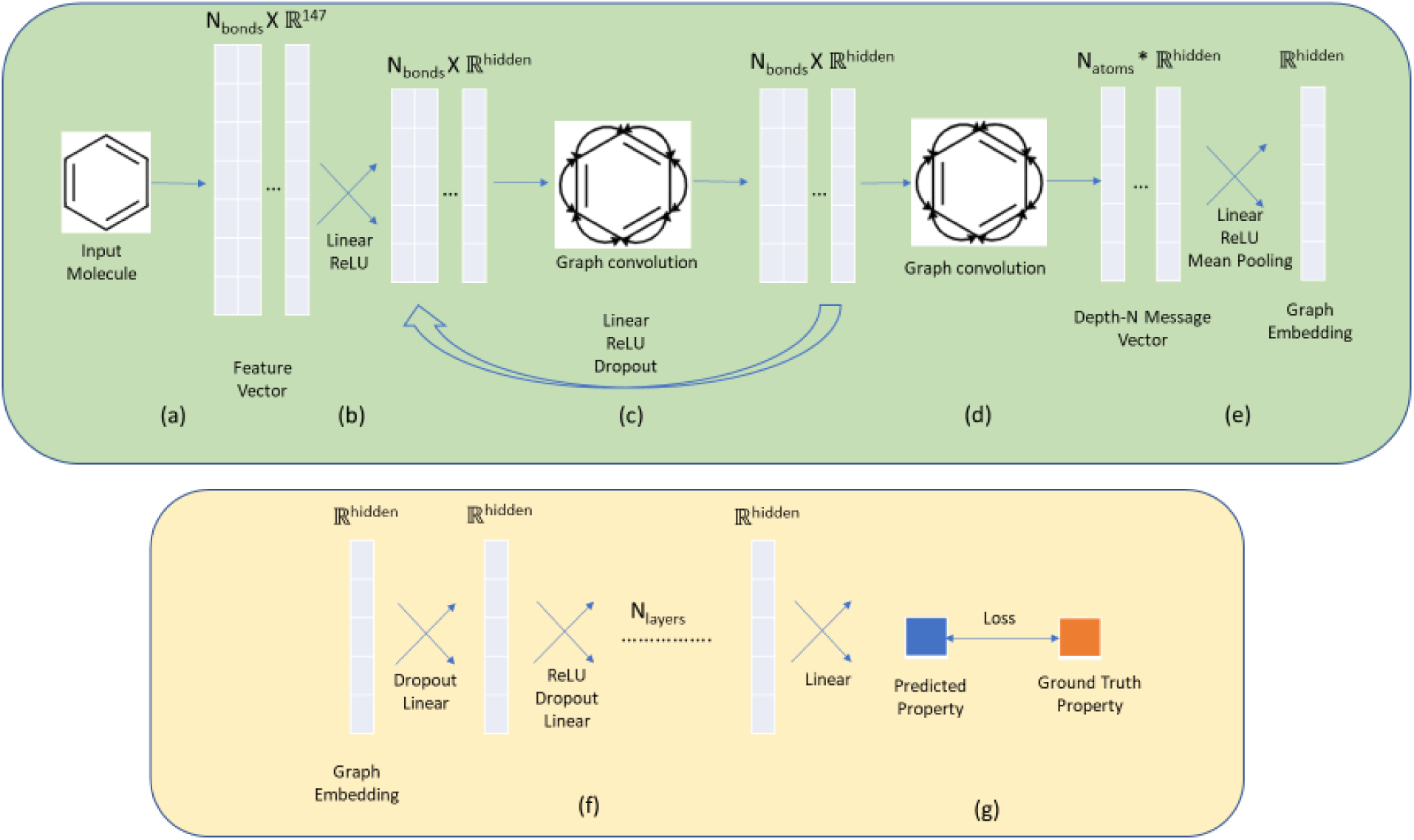
The property prediction pipeline for our method. The steps in green represent the feature extraction using Graph Convolution and the steps in orange represent regression of property scores. (a) The molecule is represented is a feature vector with features described as in section 2.1. (b) The feature vector is passed through a linear layer to get Depth-0 message. (c) Through repeated graph convolution (message passing) followed by Linear Layer, we get Depth N-1 message. (d) Each atom’s final message is calculated by summing up the messages (also Graph Convolution) of the neighbouring atoms. (e) The resultant message is passed through a Linear Layer and the mean of all the atoms is taken to get the final embedding. (f) The property score is regressed from the graph embedding by a Feed Forward Neural Network. (g) The loss between predicted property and ground truth property is then backpropagated to change the weights.

**Figure 3.**
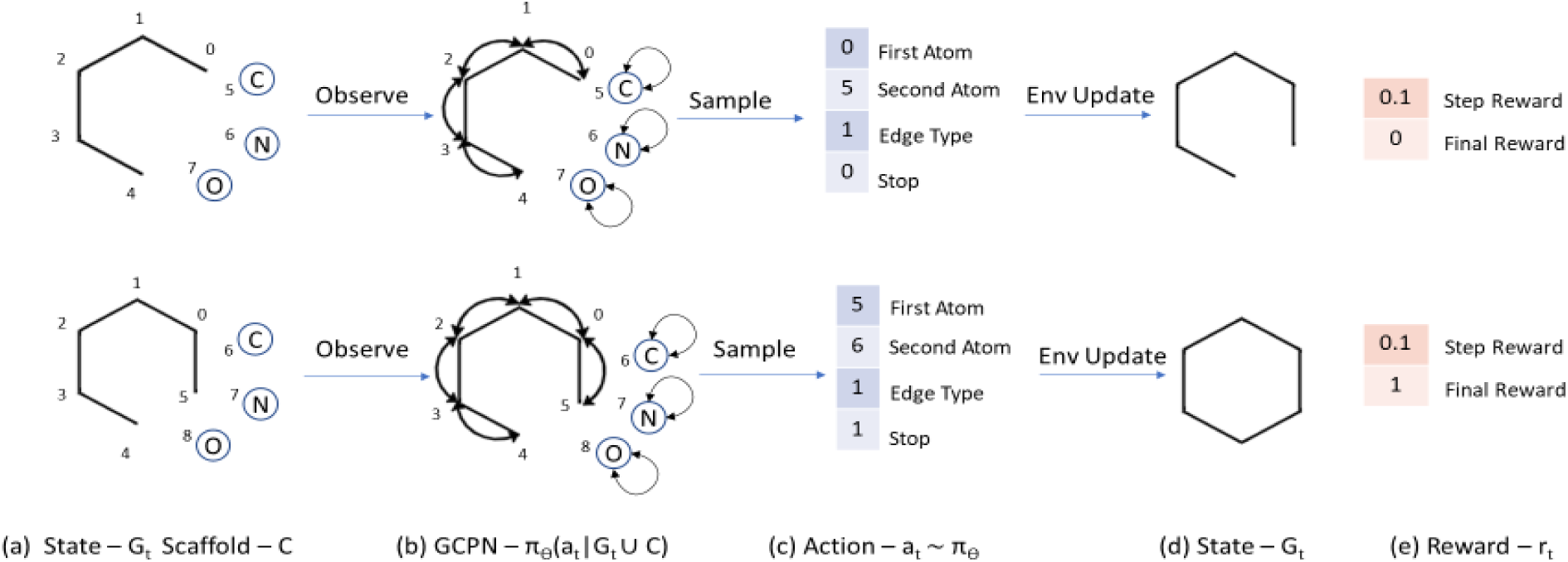
The reinforcement learning pathway for systemic generation of molecules (Redrawn from You *et al. [34]*). (a) The state is defined as the current graph *G*_*t*_ and the possible atom types *C*. (b) The GCPN conducts message passing to encode the state as node embeddings and estimates the policy function. (c) The action to be performed (*a*_*t*_) is sampled from the policy function. The environment performs a chemical valency check on the intermediate state and returns (d) the next state *G*_*t*_ and (e) the associated reward (*r*_*t*_).

**Figure 1:**
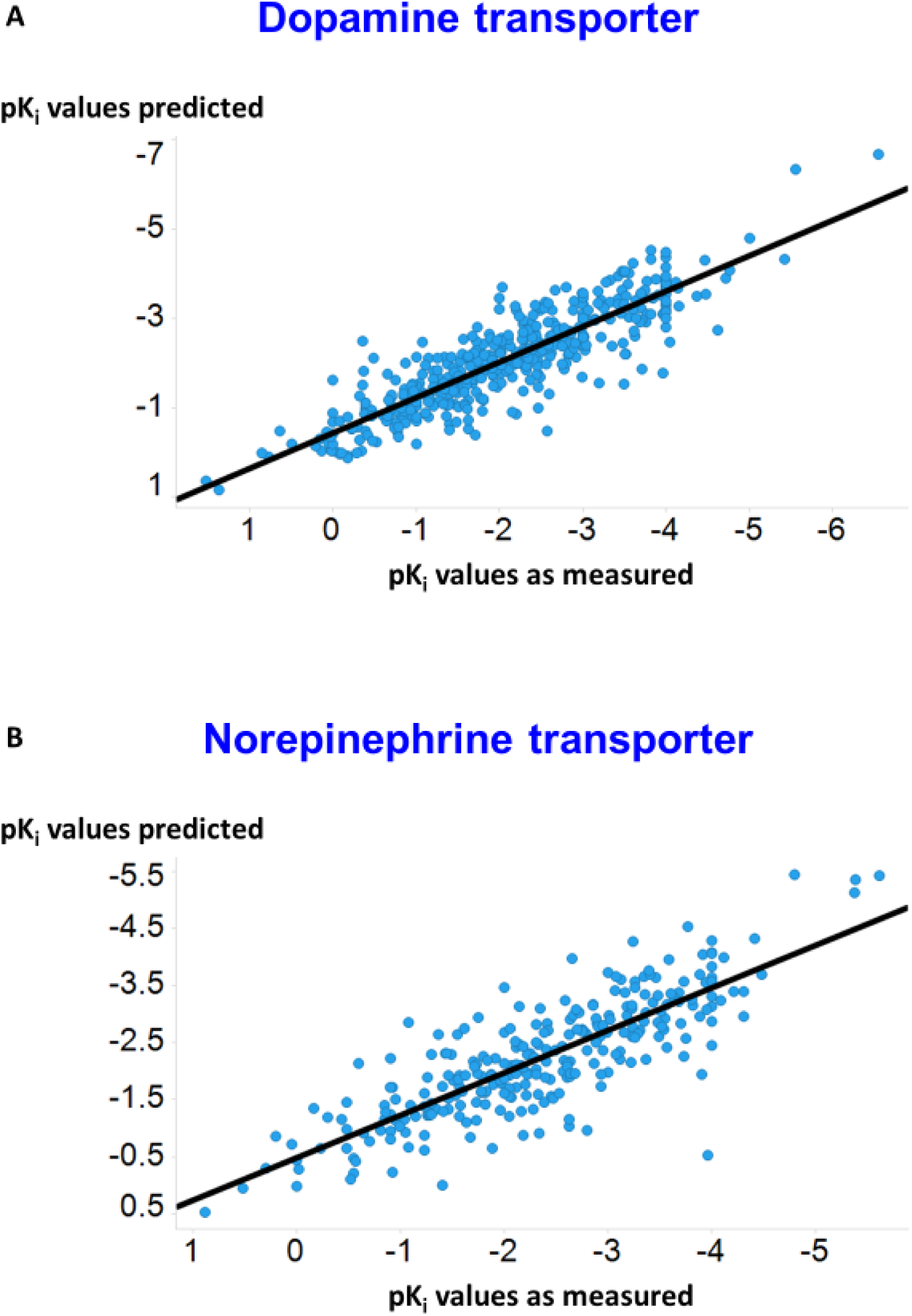
Predicted and experimental values for the test sets of the dopamine (**A**) and norepinephrine (**B**) transporters. Lines are lines of best fit (**A**: y =0.44 + 0.79x, r^2^ = 0.79; **B**: y = 0.49 + 0.74x, r^2^ =0.68).

#### 3.1.1 Hyperparameter Optimization

As the property prediction is a general algorithm with a large number of hyperparameters, we attempted to improve generalisation on the transporter problem using Bayesian optimization on the RMSE error between the predicted pKi values and the actual pKi values of the validation set. For this task we consider the hyperparameters to be the depth of the GCN encoder, the dimensions of the message vectors, the number of layers in the Feed Forward Network, and the Dropout constant.

For the case of the dopamine transporter, the optimum hyperparameters that were obtained are 3 (depth of GCN), 1300 (dimensions of message vector), 2 (FFN layers) and 0.1 (Dropout).The RMS error on the test dataset for the dopamine transporter after Hyperparameter Optimization was found to be 0.57 as compared to an error of 0.65 without it. We attribute this quite significant remaining error to the errors present in the dataset. Similarly for the norepinephrine transporter, the test RMS error was found to be 0.66 after hyperparameter optimization and the optimum hyperparameters obtained are 5 (depth of GCN), 900 (dimensions of message vector), 3 (FFN layers), 0.15 (Dropout).

#### 3.1.2 Implementation details

For the prediction of pKi value of both Dopamine and Norepinephrine transporters, we split the overall dataset into train (80%), validation (10%) and test (10%) datasets randomly. The training is done with a batch size of 50 molecules and for 100 epochs. All the network weights were initialized using Xavier initialization [75]. The first two epochs are warmup epochs [76] where the learning rate increases from 1e-4 to 1e-3 linearly and after that it decreases exponentially to 1e-4 by the last epoch. The code was written using the PyTorch library and the training was done using an NVIDIA RTX 2080Ti GPU on a Windows 10 system with 256GB RAM and an Intel 18-Core Xeon W-2195 processor.

### 3.2 Single-Objective Molecular Generation

To begin the RL evaluation we consider molecular generation with a single objective (dopamine transporter interaction). For all the experiments we use the following implementation details. The learning rate for training all the networks is taken to be 1e-3 and linearly decreasing to 0 by 3e7 timesteps. The depth of GCN network for both the GCPN and the Discriminator network is taken to be 3 and the node embedding size was taken to be 128. The code was written using the TensorFlow library and training was done using an NVIDIA RTX 2080Ti GPU

For the task of analysing the results we provide the ‘top 10’ molecules generated as in Fig 5. However, we aim to generate molecules that are in some sense similar to the original training dataset by systematically modifying the RL pathway in the following experiments. For each experiment, we find the closest molecule in the bindingDB dataset to the top 10 generated molecules. The relative closeness is measured by calculating its Tanimoto Similarity between the RDKit fingerprints, and we visualize the distribution of the TS values.

**Figure 5.**
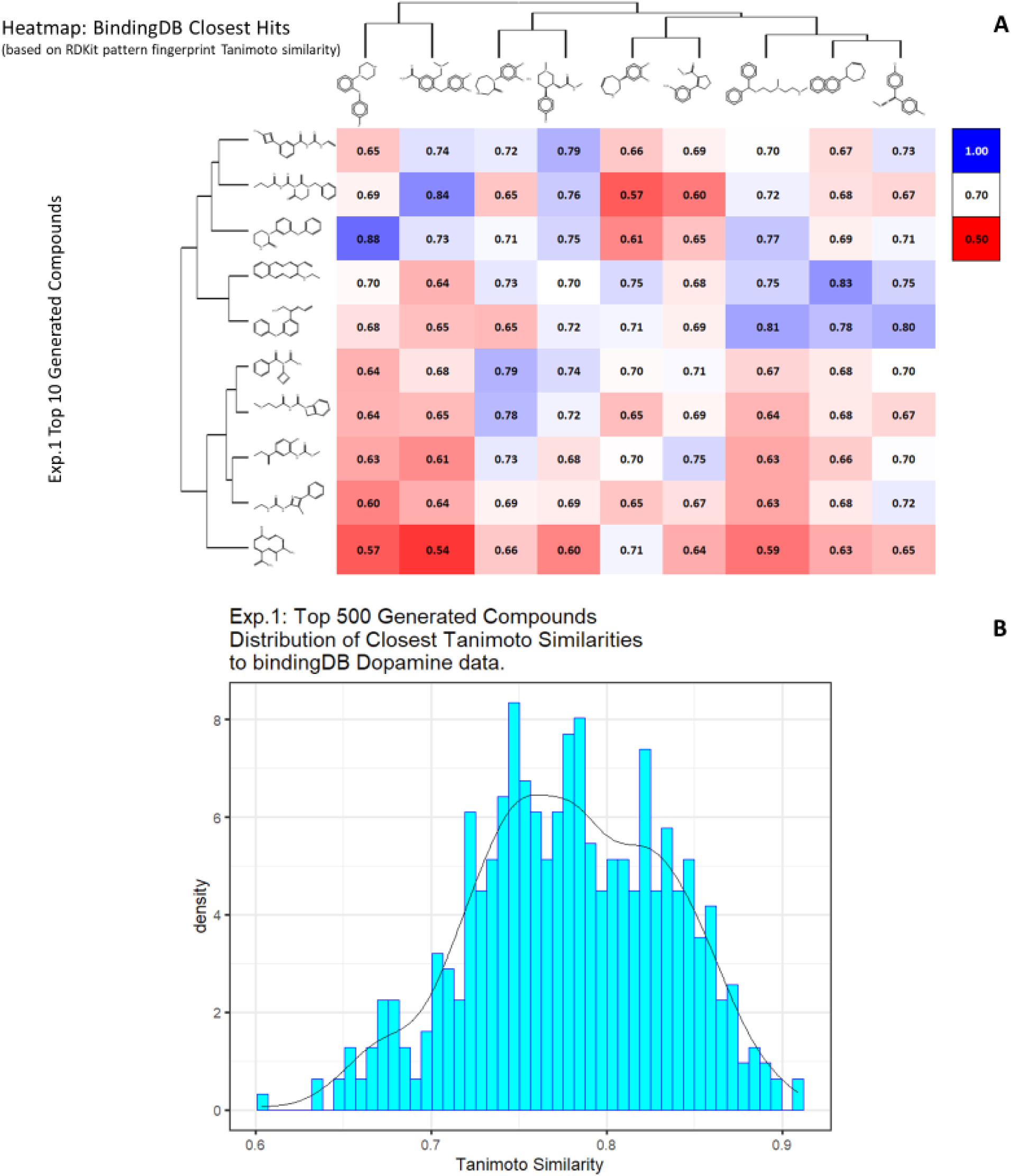
*In silico* generation by DeepGraphMol of novel molecules with predicted binding capacity to the dopamine transporter. Molecules were generated as described in the text. **A**. Top 10 molecules as predicted by DeepGraphMol versus the closest molecule in the BindingdB dataset and the Tanimoto similarity thereto (encoded using the RDKit patterned fingerprint). **B**. Distribution of Tanimoto similarities to a molecule in BindingdB dataset of the top 500 molecules.

First, we initialize the molecule with a single Carbon atom in the beginning of the generative process. The expert dataset in this case is chosen to be the ZINC dataset [77], which is a free dataset containing (at that time) some 230M commercially available compounds. However, for our experiments, we use 250K randomly selected molecules from ZINC as our expert dataset to make the experiments computationally tractable. The top generated molecules and their predicted properties are given in Supplementary Table 1 (including data on QED and SA) with a subset of the data illustrated in Fig 5. Note that in all cases the values of QED and SA both exceeded 0.8.

Although the above experiment was able to generate optimized molecules, there is no certainty that the predictions are correct due to the errors in the model as well as the errors that were propagated by the experimental errors in the data. We thus attempt to generate molecules that are similar to the more potent molecules. In the next experiment, we choose the expert dataset to be the original dataset on which we trained the molecules (we will call this the Dopamine Dataset), while omitting molecules having K_i_ greater than 1000. We again choose the initial molecule to be a single carbon atom. The equivalent data are given in Supplementary Table 2, with similar plots to those of Fig 5 given in Fig 6.

**Figure 6.**
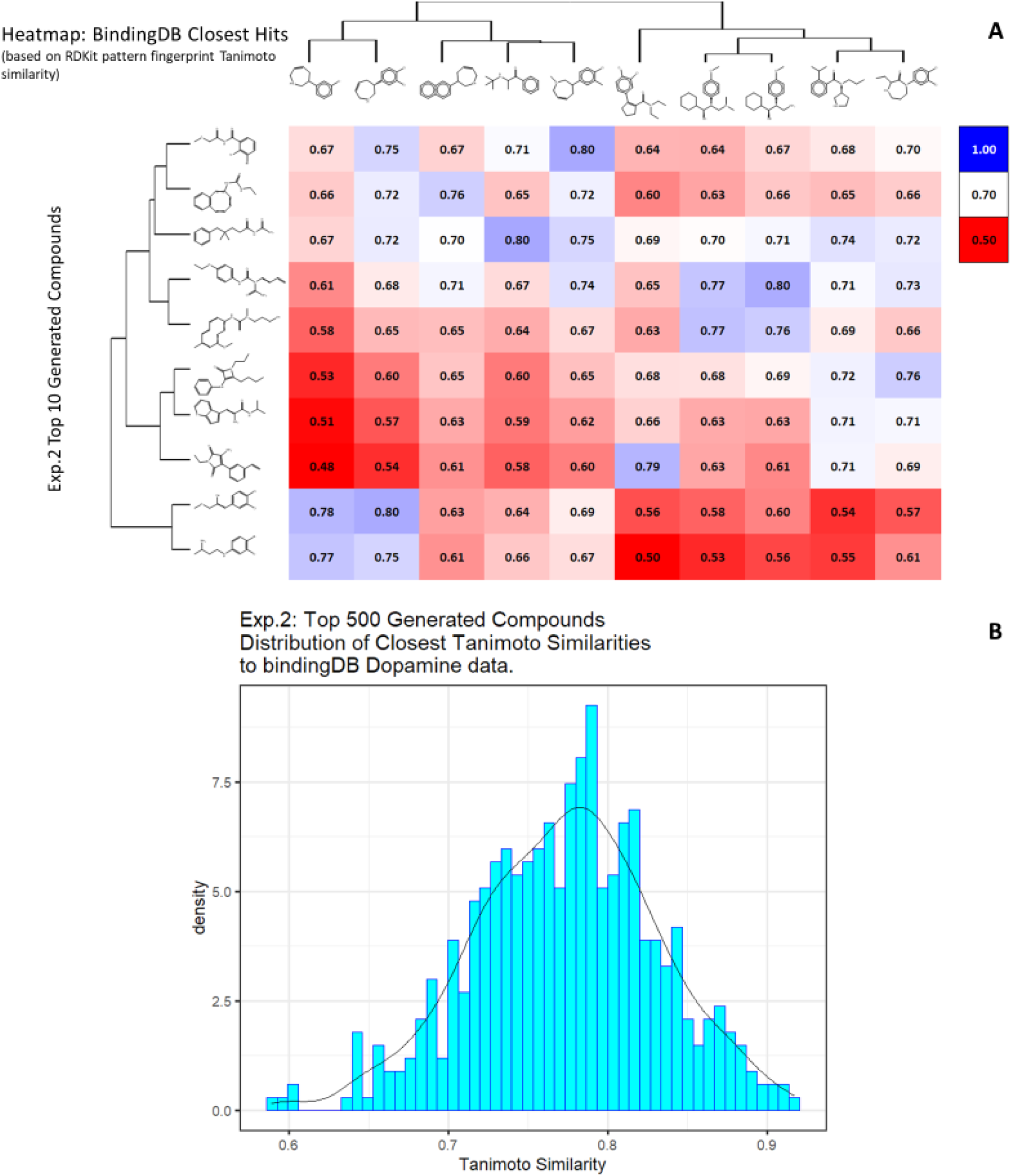
*In silico* generation by DeepGraphMol of novel molecules with predicted binding capacity to the dopamine transporter. Molecules were generated as described in the text. **A**. Top 10 molecules as predicted by DeepGraphMol versus the closest molecule in the BindingdB dataset and the Tanimoto similarity thereto (encoded using the RDKit patterned fingerprint). **B**. Distribution of Tanimoto similarities to a molecule in BindingdB dataset of the top 500 molecules.

Another way to ensure that the generated molecules will have a high affinity towards dopamine transporter is to explicitly ensure that the molecules have higher TS with already known molecules that have high pK_i_ values. We attempt to achieve this by initializing the generative process with a random molecule from the Dopamine Dataset having Ki < 1000. We conduct two experiments using this process, one where we restrict the number of atoms (other than hydrogen) to be lower than 25 (Supplementary Table 3 and Fig 7), and another (Supplementary Table 4 and Fig 8) where we restrict the number of atoms to be less than 15. For both these experiments, we use the ZINC dataset as the expert dataset. The results are summarized in the tables below. Note that in some cases we obtain a TS of 1; this is encouraging as in this case the algorithm found no need to add anything to the original molecule and could recapitulate it.

**Figure 7.**
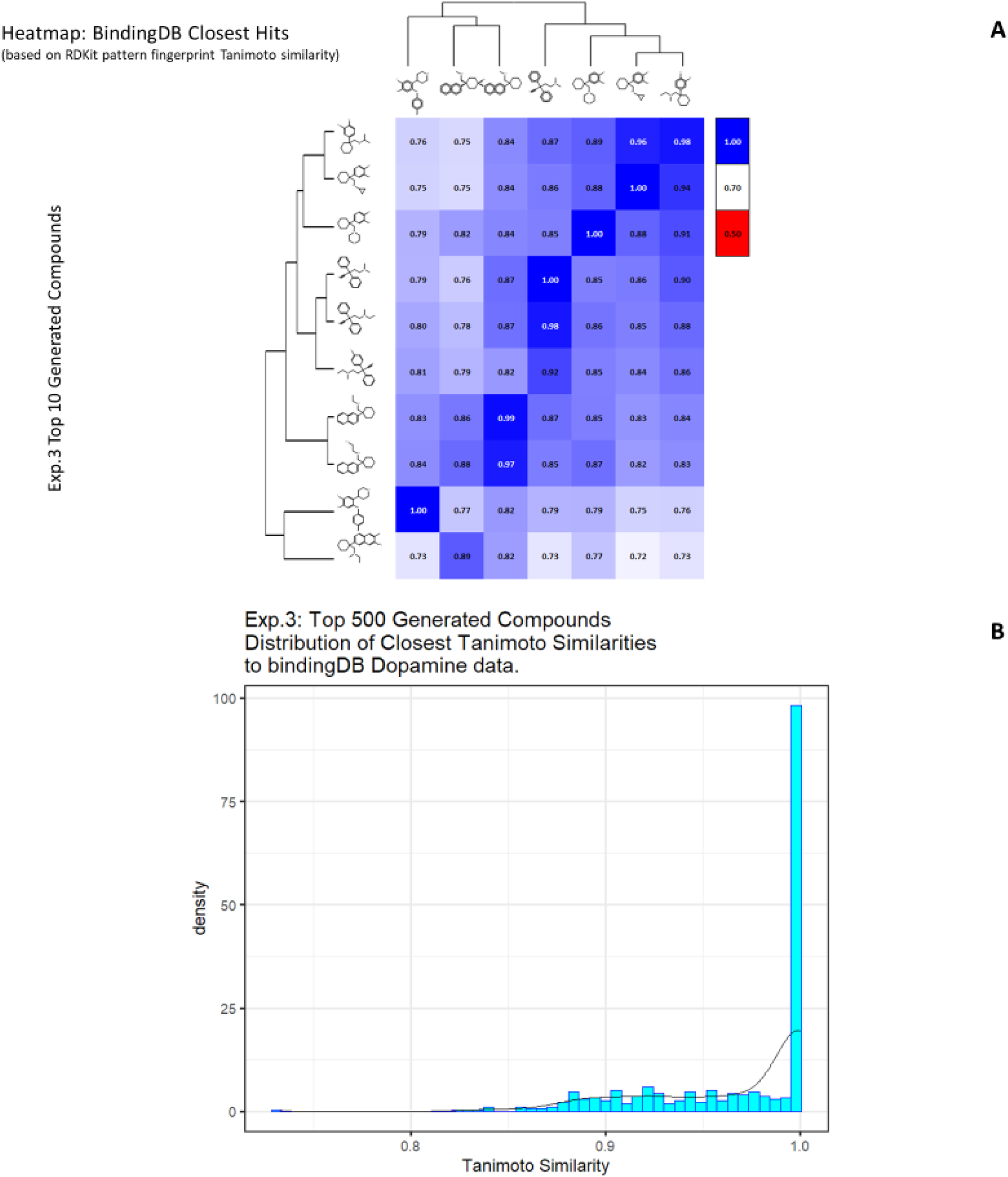
*In silico* generation by DeepGraphMol of novel molecules with predicted binding capacity to the dopamine transporter using a generative method in which the number of heavy atoms is constrained to be lower than 25. Molecules were generated as described in the text. **A**. Top 10 molecules as predicted by DeepGraphMol versus the closest molecule in the BindingdB dataset and the TS thereto (encoded using the RDKit patterned fingerprint). **B**. Distribution of Tanimoto similarities (RDKit patterned encoding) to a molecule in BindingdB dataset of the top 500 molecules.

**Figure 8.**
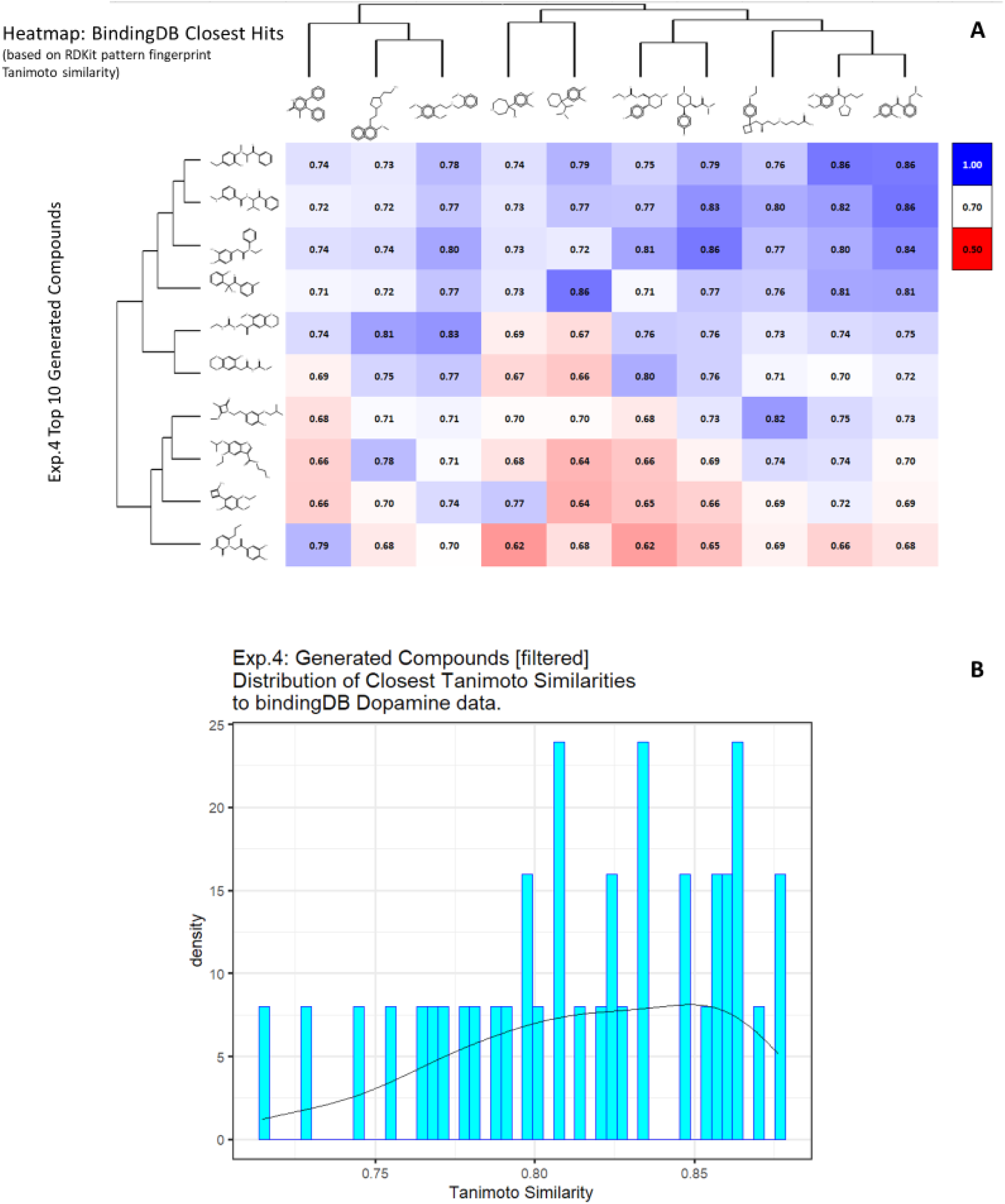
*In silico* generation by DeepGraphMol of novel molecules with predicted binding capacity to the dopamine transporter using a generative method in which the number of heavy atoms is constrained to be lower than 15. Molecules were generated as described in the text. **A**. Top 10 molecules as predicted by DeepGraphMol versus the closest molecule in the BindingdB dataset and the TS thereto (encoded using the RDKit patterned fingerprint). **B**. Distribution of Tanimoto similarities (RDKit patterned encoding) to the closest molecule in BindingdB dataset of the top 500 molecules.

### 3.3 Multi-Objective Molecular Generation

Even though generating molecules having higher affinity towards a particular ligand in itself is quite sought after, in many cases we might wish to seek molecules that bind to one receptor but explicitly do not bind to another one (kinase inhibitors might be one such example). We attempt to achieve this here with the help of our Reinforcement Learning pipeline by modifying the reward function to be a weighted combination of pK_i_ values for the two different targets. Explicitly, we attempt to generate molecules that have high binding affinity to the Dopamine Transporter but a much lower binding affinity to the Norepinephrine Transporter. Thus, we modify the reward function used in the previous experiments to add 2 times the predicted pK_i_ values for Dopamine Transporter and -1 times the predicted pK_i_ values for the Norepinephrine Transporter. The higher weight is given to the dopamine component since we wish to generate molecules that do bind to it. Clearly we could use any other weightings as part of the reward function, so those chosen are simply illustrative. For this experiment we initialize the process with a random molecule from the Dopamine dataset having a number of atoms lower than 25 and choose the expert dataset to be ZINC. The results of this experiment are summarized in Supplementary Table 5 and Fig 9. As above, some molecules have a TS of 1 in the dataset, for the same reasons.

**Figure 9.**
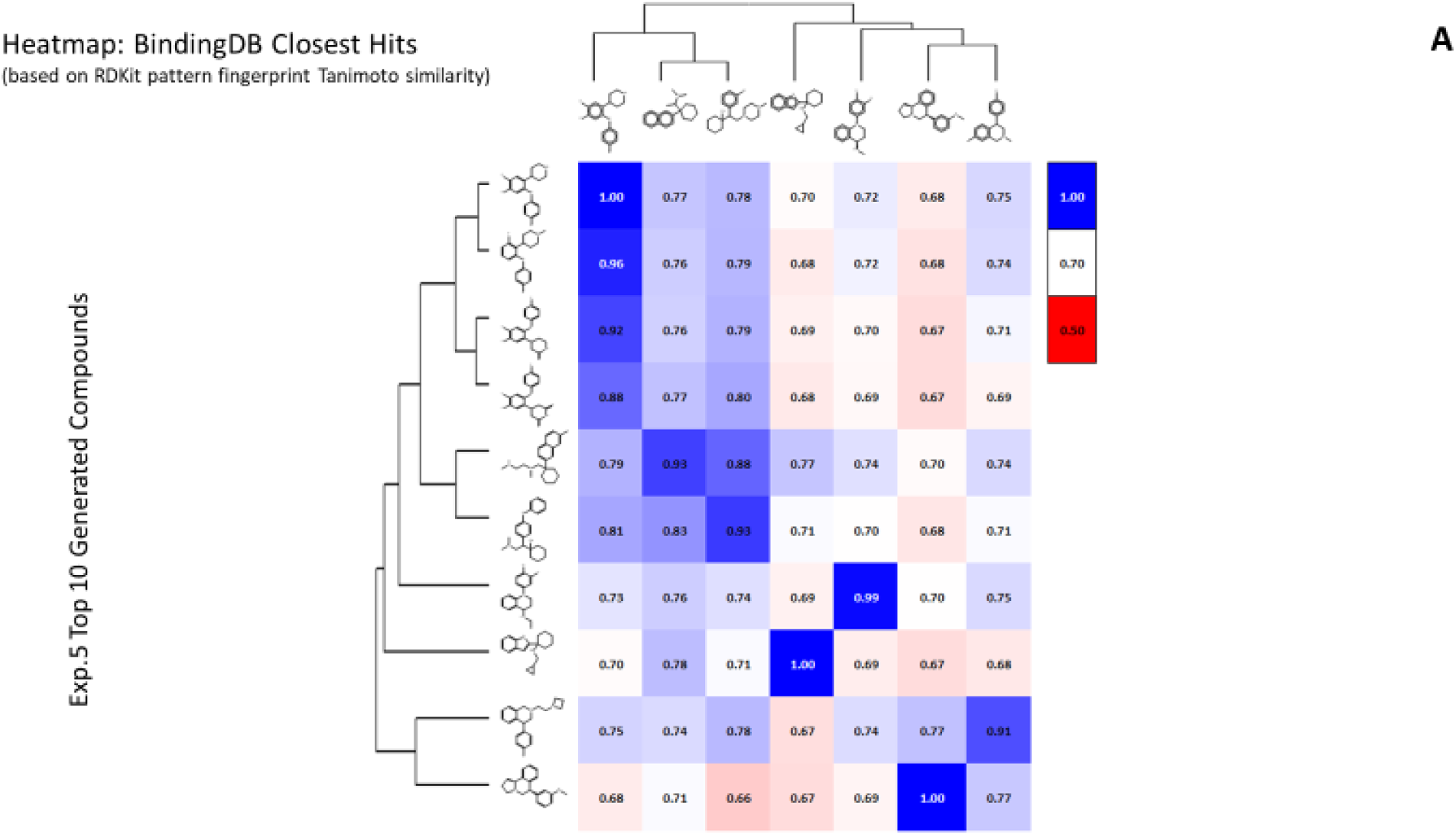
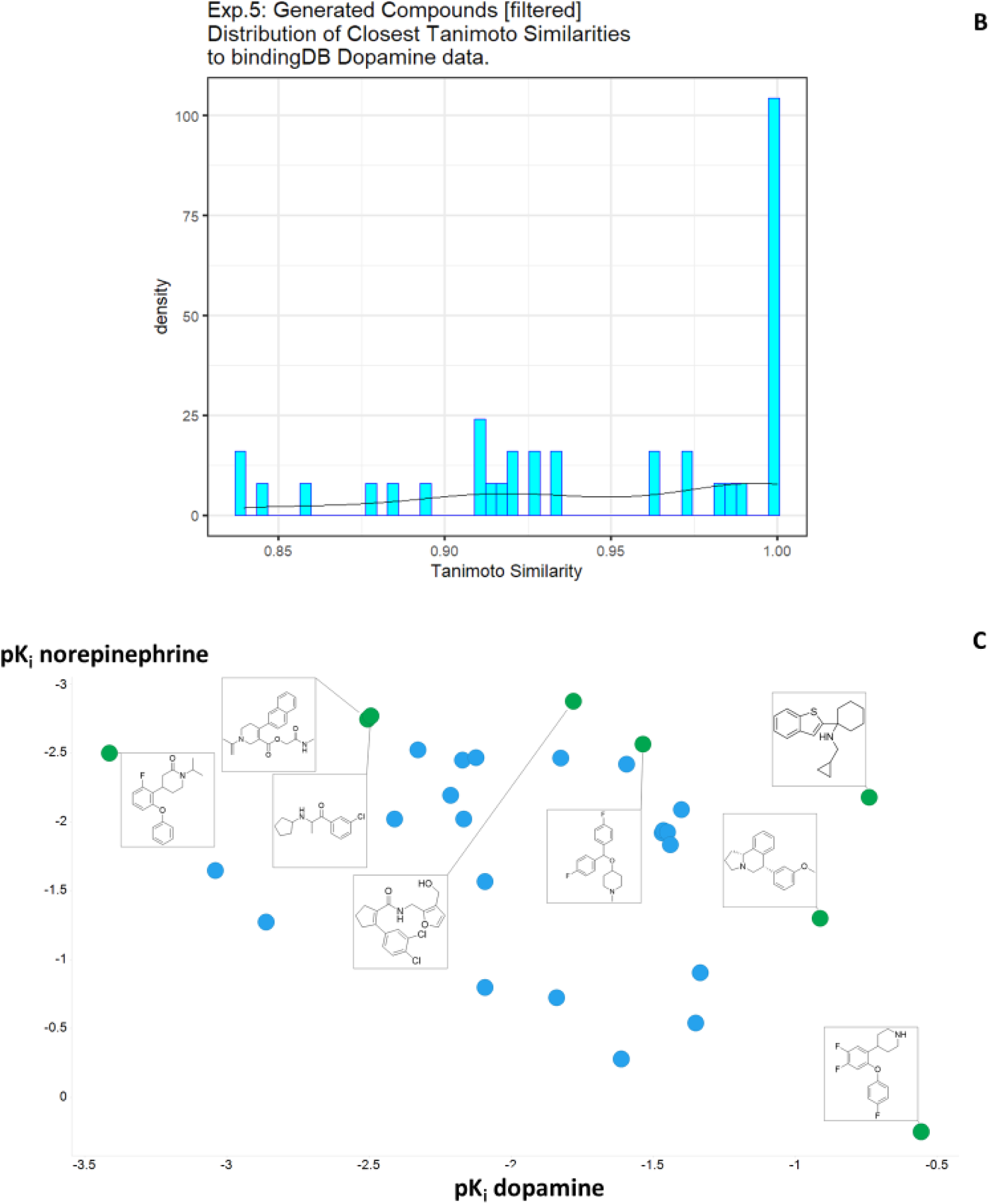
*In silico* generation by DeepGraphMol of novel molecules with predicted binding capacity to the dopamine transporter using a generative method in which the number of heavy atoms is constrained to be lower than 25. Molecules were generated as described in the text. **A**. Top 10 molecules as predicted by DeepGraphMol versus the closest molecule in the BindingdB dataset and the TS thereto (encoded using the RDKit patterned fingerprint). **B**. Distribution of Tanimoto similarities (RDKit patterned encoding) to the closest molecule in BindingdB dataset of the top 500 molecules. **C**. Plot of those molecules with differential affinities for the dopamine and norepinephrine transporters.

Only in rare cases do candidate solutions for multi-(in this case two-)objective optimisation problems have unique solutions that are optimal for both [78], and there is a trade-off that is left to the choice of the experimenter. Thus, Fig 9C also illustrates the molecules on the Pareto front for the two objectives, showing how quite changes in structure can move one swiftly along the Pareto front. Consequently our method also provides a convenient means of attacking multiobjective molecular optimisation problems.

## Conclusions

Overall, the present molecular graph-based generative method has a number of advantages over grammar-based encodings, in particular that it necessarily creates valid molecules. As stressed by Coley and colleagues [49], such methods still retain any inherent limitations of 2D methods as *a priori* they do not encode 3D information. This said, there is evidence that 3D structures do not add much benefit when forming QSAR models [79-83], so we do not consider this a major limitation for now. Some of the molecules generated might be seen by some (however subjectively) as ‘unusual, even though they scored well on both drug-likeness and synthetic accessibility metrics. This probably says much about the size of plausible drug space that exists relative to the fraction that has actually been explored [84-86], and implies that generative methods can have an important role to play in medicinal chemistry. In conclusion, we here add to the list of useful, generative molecular methods for virtual screening by combining molecular graph encoding, reinforcement learning and multiobjective optimisation within a single strategy.

## Supporting information

Exp 1

## Acknowledgments

The work of SS and DBK is supported as part of EPSRC grant EP/S004963/1 (SuSCoRD).

